# Repurposed drugs and their combinations prevent morbidity-inducing dermonecrosis caused by diverse cytotoxic snake venoms

**DOI:** 10.1101/2022.05.20.492855

**Authors:** Steven R. Hall, Sean A. Rasmussen, Edouard Crittenden, Charlotte A. Dawson, Keirah E. Bartlett, Adam P. Westhorpe, Laura-Oana Albulescu, Jeroen Kool, José María Gutiérrez, Nicholas R. Casewell

## Abstract

Morbidity from snakebite envenoming affects approximately 400,000 people annually. Tissue damage at the bite-site often leaves victims with catastrophic life-long injuries and is largely untreatable by currently available antivenoms. Repurposing small molecule drugs that inhibit specific snake venom toxins offers a potential new treatment strategy for tackling this neglected tropical disease. Using human skin cell assays as an initial model for snakebite-induced dermonecrosis, we show that the repurposed drugs 2,3-dimercapto-1-propanesulfonic acid (DMPS), marimastat, and varespladib, alone or in combination, reduce the cytotoxic potency of a broad range of medically important snake venoms up to 5.7-fold. Thereafter, using a preclinical murine model of dermonecrosis, we demonstrate that the dual therapeutic combinations of DMPS or marimastat with varespladib potently inhibit the dermonecrotic activity of three geographically distinct and medically important snake venoms. These findings strongly support the future translation of repurposed drug combinations as broad-spectrum therapeutics for preventing morbidity caused by snakebite.

## Introduction

Current estimates suggest that 1.8-2.7 million people are envenomed due to snakebite every year, resulting in 81,000-138,000 deaths and 400,000 cases of morbidity annually, predominantly affecting those in the tropics and sub-tropics^1–3^. Snakebite has been labelled ‘the most neglected of neglected tropical diseases (NTDs)’^4^, with the late UN Secretary General Kofi Annan calling it ‘the biggest public health crisis you’ve never heard of’^5^. In 2017, snakebite envenoming was added to the World Health Organization (WHO)’s formal list of NTDs; the WHO has since elevated snakebite to a ‘priority category A NTD’ and has created a roadmap with the goal of reducing the global burden of snakebite by one-half by the year 2030^6^. One of the proposed methods to accomplish this is to develop novel treatments for snakebite; an ambitious task considering the myriad issues associated with developing snakebite therapies, including the variability and complexity of toxins that make up different snake venoms^7,8^.

Snake venoms are comprised of dozens of different toxins at varying concentrations, which differ both inter- and intra-specifically and induce a range of pathological and pathophysiological effects^7^. However, there are four primary toxin families that are dominant across many different venoms and thus represent attractive targets for toxin-inhibiting therapeutics: phospholipases A_2_ (PLA_2_s), snake venom metalloproteinases (SVMPs), snake venom serine proteases (SVSPs), and three-finger toxins (3FTxs)^9^. The main syndromes of snakebite envenoming are generally categorised as haemotoxic (e.g. haemorrhage and coagulopathy), neurotoxic (e.g. muscle paralysis), and/or cytotoxic (e.g. local tissue necrosis)^10,11^. Haemotoxicity is a particularly common sign of envenoming, especially following bites from viperid (family Viperidae) snakes, and is largely caused by SVMPs, SVSPs, and PLA_2_s^10–12^. Neurotoxic envenoming is more commonly caused by elapid (family Elapidae) snakes and is primarily associated with neurotoxic 3FTxs and PLA_2_s^11,13^. While haemotoxicity and neurotoxicity are primarily responsible for snakebite-induced mortality, life-altering morbidity in survivors is most frequently caused by severe local tissue damage in and around the bite-site. This pathology is caused by cytotoxic 3FTxs, SVMPs, and PLA_2_s and frequently leads to permanent disability, often requiring surgical debridement or even amputations of the affected limb or digit^14,15^.

The only treatments currently available for snakebite envenoming are animal-derived polyclonal antibody therapies called antivenoms. These therapies have conceptually remained unchanged for over a century and are associated with a multitude of issues including high cost, requirement for a consistent cold-chain, limited cross-snake species efficacy due to venom variation, and high frequency of adverse events post-administration^1,7,8,16–19^. In addition, due to the large size of antivenom antibodies or their fragments (i.e. typically ∼110 kDa, F(ab’)_2_; or ∼150 kDa, IgG) these treatments are unable to efficiently penetrate into peripheral tissue surrounding a bite-site thus reducing their efficacy against local tissue cytotoxicity, and they need to be administered intravenously (IV) in a clinical environment by a medical professional which severely restricts their utility in rural communities where snakebite victims are often hours or even days away from appropriate facilities^1,8,20^. To address some of these challenges, next-generation snakebite therapies such as toxin-specific monoclonal antibodies^21,22^ and toxin-inhibiting small molecule drugs^23–29^, have received considerable attention in recent years.

Small molecule drugs (hereafter simply called drugs) offer many desirable characteristics in comparison to existing antivenoms, such as increased cross-species efficacy, tolerability, stability, and affordability^8,26,27,29^. Additionally, small molecule drugs are more able to penetrate peripheral tissue than large IgG-derived antibodies, meaning they should exhibit enhanced tissue distribution dynamics to better inhibit cytotoxins at the bite site. Further, they can be formulated as oral or topical therapies which could be administered in the field much more quickly after a snakebite victim is envenomed compared to an IV-administered antivenom^8,25,26,29–32^.

Three repurposed drugs initially developed for other conditions^26,33,34^ have shown particular promise as potential new drug therapies for snakebite envenoming based on *in vitro* and rodent *in vivo* data: the SVMP-inhibiting metal chelator, DMPS (Unithiol)^26^, the hydroxamic acid, marimastat^27,28,35–37^, and the secretory PLA_2_-inhibiting drug, varespladib^23,28,38–41^. Additionally, it has been shown that combining marimastat with varespladib improves their pan-geographic utility, resulting in superior prevention of venom-induced lethality in mice compared with either drug alone against diverse snake venoms^27^. While these studies have demonstrated such drugs can effectively protect against snake venom-induced lethality in animal models, there is limited published evidence of their efficacy or potential utility against the pathology most likely to cause the permanent, life-changing injuries often seen in snakebite survivors: tissue necrosis caused by snake venom cytotoxins. Herein we explore the therapeutic potential of small molecules drugs against the local tissue damage stimulated by cytotoxic snakebite envenoming. Using a variety of geographically diverse snake venoms, we demonstrate that DMPS, marimastat, and varespladib individually provide protection against snake venom cytotoxins to different extents, but that drug combinations are highly effective at preventing local tissue damage *in vivo*, and thus represent promising leads for combatting the local dermonecrotic effects caused by snakebite envenoming.

## Results

### Diverse snake venoms inhibit human epidermal keratinocyte viability

Prior to exploring the inhibitory capability of drugs against the cytotoxic effects of snake venoms, we first defined the effect of 11 venoms sourced from distinct snake species and geographic regions on the viability of adherent human skin cells. Using 3-(4,5-dimethylthiazol-2-yl)-2,5-diphenyl tetrazolium bromide (MTT) assays^42,43^ and immortalised human epidermal keratinocytes (HaCaT^44,45^), we generated venom dose-response curves (**Fig. 1a-k**). MTT assays measure two types of venom action on adherent cells: direct inhibition of cell viability^42,43^ and cellular detachment from the culture plate (an effect that can be caused by certain SVMPs, such as BAH1^46^), both of which evidence the deleterious actions of venoms on the keratinocytes. Using a broad concentration range for each venom and measuring the resulting viability of adherent cells after 24 hours, we calculated the concentration at which cell viability was inhibited for each venom by 50% (IC_50_ values; **Fig. 1l**) as a measure of potency. Our results demonstrated that 9 of the 11 venoms tested displayed similar potencies, with those from the vipers *Bitis arietans* (puff adder, sub-Saharan Africa), *Bothrops asper* (fer-de-lance, Central America), *Crotalus atrox* (Western diamondback rattlesnake, North America), *Calloselasma rhodostoma* (Malayan pit viper, South East Asia), *Echis carinatus* (Indian saw-scaled viper, South Asia and the Middle East) and *Echis ocellatus* (West African carpet viper, West Africa) (IC_50_ range: 7.5 – 19.6 μg/mL) comparable to those from the elapid spitting cobras *Naja nigricollis* (black necked spitting cobra, West Africa and East Africa variants) and *Naja pallida* (red spitting cobra, East Africa) (IC_50_ range: 23.1 – 27.2 μg/mL). The venom of *Daboia russelii* (Russell’s viper, South Asia; IC_50_: 45.1 μg/mL) was slightly, albeit significantly, less potent than that of the other vipers *B. asper, C. atrox, C. rhodostoma, E. carinatus*, and *E. ocellatus*, while venom from the primarily neurotoxic non-spitting cobra, *Naja haje* (Egyptian cobra) (IC_50_: 86.8 μg/mL), was the least potent with a significantly higher IC_50_ value than all other tested venoms. None of the resulting Hill slopes, measures of the steepness of each venom’s dose-response curve, were significantly different from each other, though all 11 were greater than |-1| (**Fig. 1m**), suggesting likely ‘positive cooperativity’ between venoms toxins^47,48^.

**Fig. 1.**
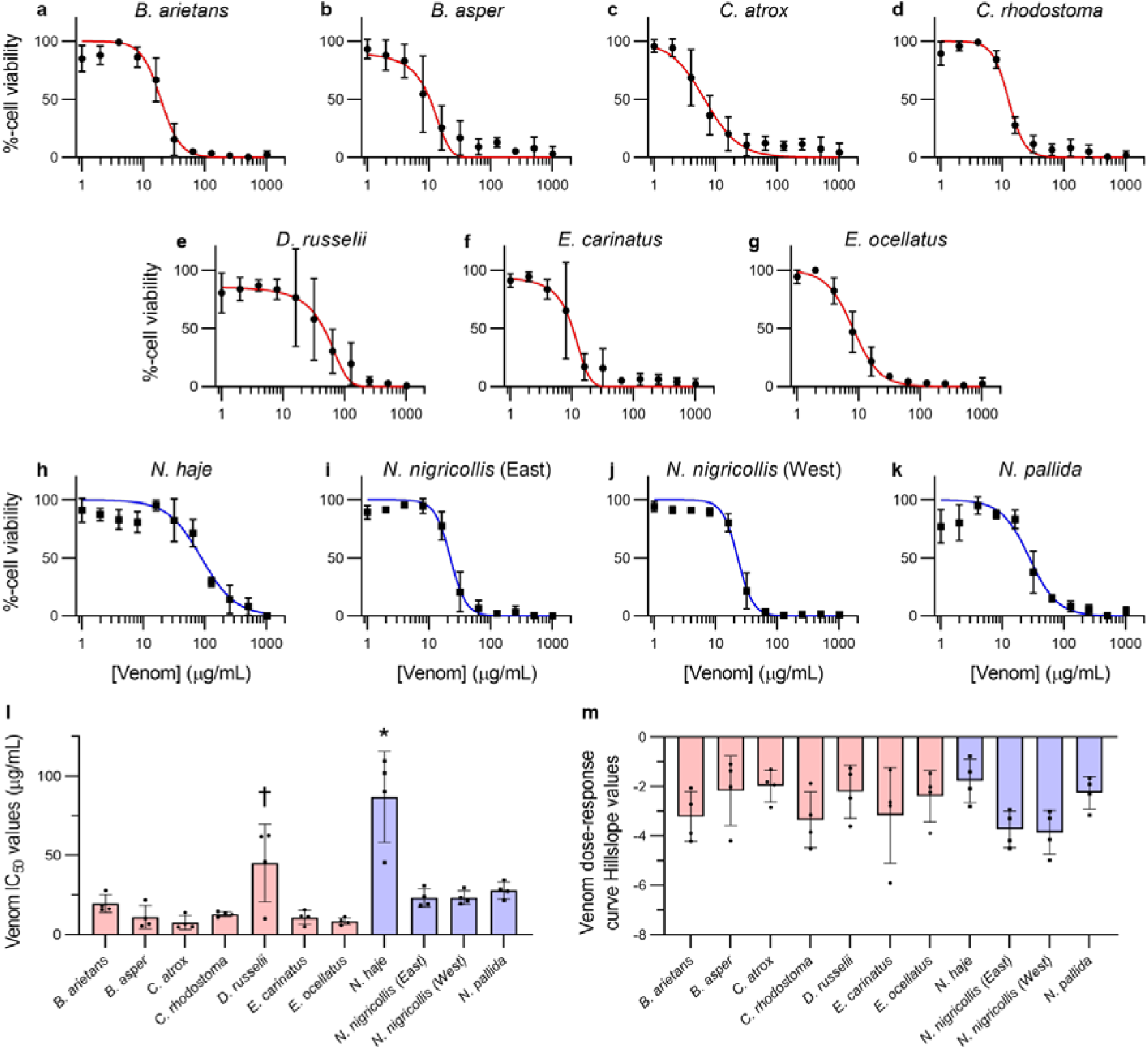
Snake venoms dose-dependently inhibit HaCaT adherent cell viability. MTT cell viability assays were completed in adherent HaCaT epidermal keratinocytes exposed to serial dilutions (1 – 1,024 μg/mL) of different snake venoms for 24 hours. The venoms tested were from (**a**) *Bitis arietans*, (**b**) *Bothrops asper*, (**c**) *Crotalus atrox*, (**d**) *Calloselasma rhodostoma*, (**e**) *Daboia russelii*, (**f**) *Echis carinatus*, (**g**) *Echis ocellatus*, (**h**) *Naja haje*, (**i**) East African *Naja nigricollis*, (**j**) West African *Naja nigricollis*, and (**k**) *Naja pallida*. (**l**) IC_50_ and (**m**) Hill slope values were calculated for each independent trial. **Red-**coloured data denotes viperid snakes, while **blue**-coloured data denotes elapid snakes. * Signifies that the value is significantly higher than all other tested venoms, and † signifies that the value is significantly higher than *B. asper, C. atrox, C. rhodostoma, E. carinatus*, and *E. ocellatus*, as determined by a one-way ANOVA comparing all values to each other followed by a Tukey’s multiple comparisons test (*P* < 0.05, n=4). ANOVA statistics for individual statistically analysed graphs are: (**l**) F(10,33) = 14.47, P<0.0001; (**m**) F(10,33) = 1.828, P=0.0942. Error bars represent SD of four independent trials, and the individual IC_50_ and Hill slope values for each trial are shown as points within the bars of the graphs in panels L and M.

### DMPS and marimastat reduce the loss of adherent cell viability stimulated by certain snake venoms

Prior to investigating the inhibitory potency of toxin-inhibiting drugs in the MTT assay, we first determined the cellular ‘maximum tolerated concentration (MTC)’ of the repurposed drugs DMPS, marimastat, and varespladib. Thus, HaCaT cells were treated with two-fold serial dilutions of each drug until a significant reduction in cell viability was observed after 24 hours of exposure. The highest concentration of each drug that did not significantly reduce cell viability when compared to the vehicle control (labelled ‘0’) was determined to be the MTC. Then, to ensure that cells would be treated with a sub-toxic amount of drug in the venom-inhibition experiments, one half of this dose (MTC_½_) was selected for the venom-drug co-treatment experiments^49,50^. The MTC_½_ for DMPS, marimastat, and varespladib used in the following experiments were 625, 2.56, and 128 μM, respectively (**Supplementary Fig. 1**).

Next, using a drug pre-incubation model^26,27^ followed by MTT assays in the HaCaT cells, we tested the inhibitory effect of the three toxin-inhibiting drugs (using their MTC_½_ values) against six of our previously tested cytotoxic snake venoms. Our results demonstrated that the SVMP inhibitors DMPS and marimastat^26,27,35^ significantly (*P* < 0.05) reduced the cell-damaging potency of venom from *C. atrox, E. carinatus*, and *E. ocellatus* (**Fig. 2b-d**, respectively), as demonstrated by the increased IC_50_ values. Additionally, DMPS slightly, albeit significantly, increased the IC_50_ of East African *N. nigricollis* venom (*P* = 0.0053) (**Fig. 2e**), though its effect was not significant against West African *N. nigricollis* venom (*P =* 0.0501) (**Fig. 2f)**. In contrast, the PLA_2_-inhibitor varespladib^23^ did not display an inhibitory effect on any of the six tested venoms. The cell viability-inhibitory effects of *B. arietans* and West-African *N. nigricollis* (**Fig. 2a,f**) venom were not significantly inhibited by any of the tested drugs.

**Fig. 2.**
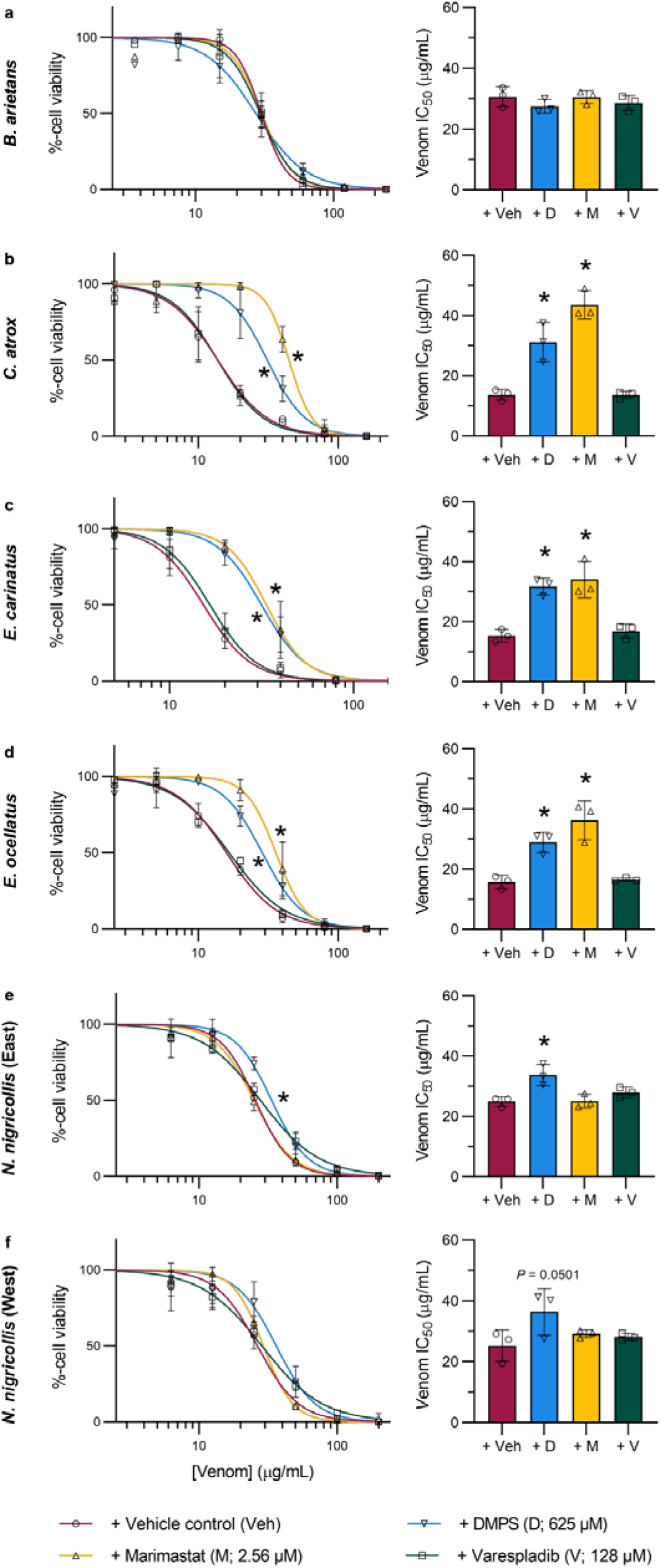
DMPS and marimastat, but not varespladib, inhibit the potency of certain cytotoxic snake venoms in adherent HaCaT cells. Serial dilutions of venoms (2.5 – 200 μg/mL) were pre-incubated with the MTC_½_ of DMPS, marimastat, varespladib, or vehicle control for 30 minutes, after which HaCaT cells were exposed to the treatments for 24 hours followed by MTT cell viability assays, from which venom concentration-response curves and their associated IC_50_ values were calculated. Panels show venom from (**a**) *B. arietans*, (**b**) *C. atrox*, (**c**) *E. carinatus*, (**d**) *E. ocellatus*, (**e**) East African *N. nigricollis*, and (**f**) West African *N. nigricollis*. * Signifies that the IC_50_ is significantly higher than that of the vehicle control for that venom as determined by a one-way ANOVA followed by Dunnett’s multiple comparisons test (*P* < 0.05, n = 3). ANOVA statistics for individual statistically analysed graphs are: (**a**) F(3,8) = 1.057, P=0.4195; (**b**) F(3,8) = 37.16, P<0.0001; (**c**) F(3,8) = 21.17, P=0.0004; (**d**) F(3,8) = 20.34, P=0.0004; (**e**) F(3,8) = 8.757, P=0.0066; (**f**) F(3,8) = 2.998, P=0.0952. Error bars represent SD of at least three independent trials, and the individual values for each trial are shown as points within each of the bar graphs.

### DMPS and marimastat, but not varespladib, inhibit PLA_2_-rich *D. russelii* and *B. asper* venoms

Due to the surprising lack of inhibitory effect observed with varespladib in the MTT cell viability studies summarised in **Fig. 2**, we decided to repeat these experiments using venoms from *D. russelii* and *B. asper*, which have higher PLA_2_ toxin abundances proportionally than the other six tested venoms^9^, and to increase the concentration of varespladib from its MTC_½_ (128 μM) to its MTC (256 μM). In addition, propidium iodide (PI) cell death assays^51,52^ were multiplexed with the MTT assays as secondary measures of the cytotoxic potencies of the venoms, in case varespladib was incompatible with the MTT assays. Despite the potential for more abundant PLA_2_ toxins to contribute to cell cytotoxic effects, varespladib again showed no inhibition as measured by either MTT or PI assays against either of these viper venoms (**Fig. 3**). None of the drugs significantly inhibited *D. russelii* venom potency as measured with MTTs, though DMPS reduced its potency as measured with PI (**Fig. 3a-b**). Both DMPS and marimastat inhibited *B. asper* venom potency as measured by MTT, while only marimastat did so as measured by PI (**Fig. 3c-d**).

**Fig. 3.**
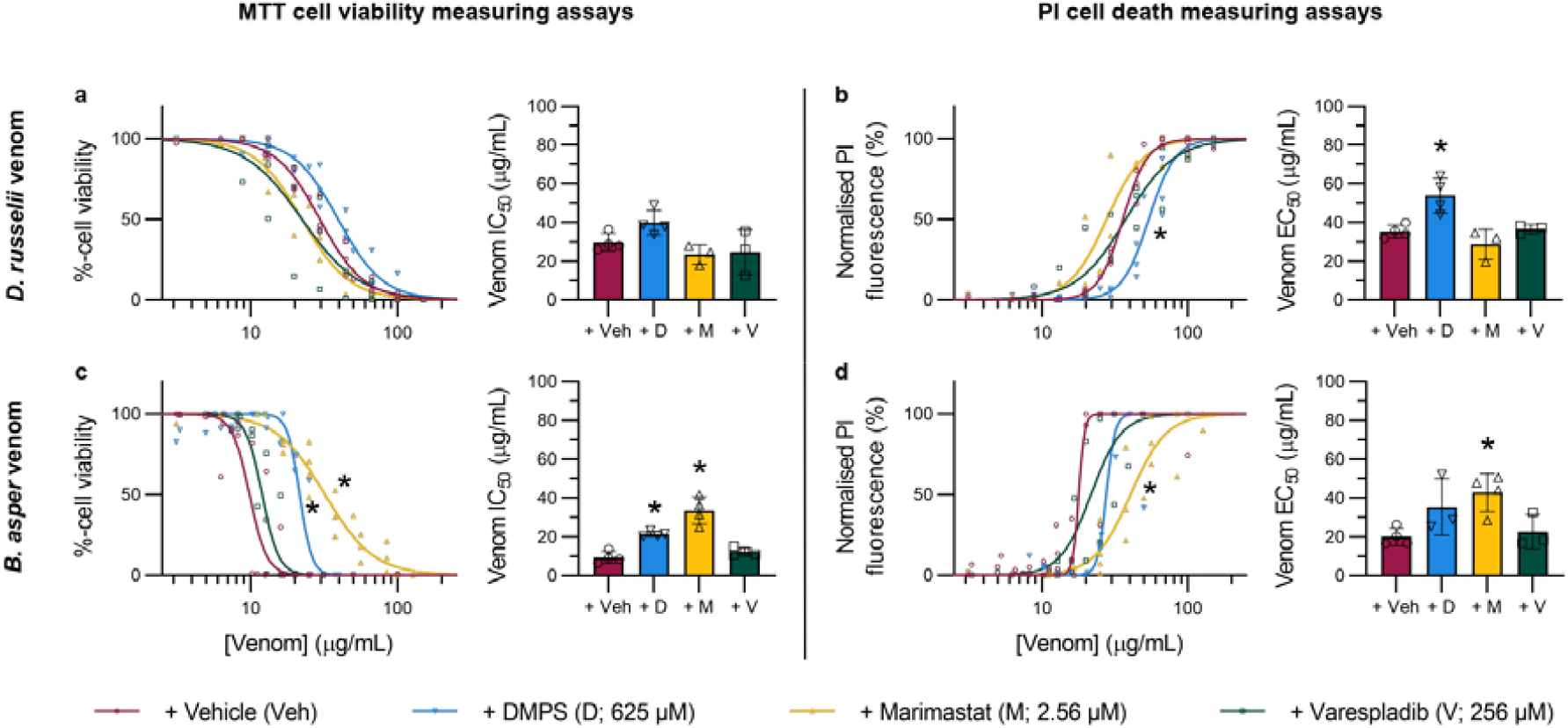
SVMP inhibitors reduce the loss of HaCaT cell viability and/or cell death stimulated by *D. russelii* and *B. asper* venoms. HaCaT cells were treated for 24 hours with serial dilutions of *D. russelii* (3.125 – 100 μg/mL, **top row**) or *B. asper* (2.2 – 127 μg/mL, **bottom row**) venom that had been pre-incubated with drug vehicle control, DMPS (625 μM), marimastat (2.56 μM), or varespladib (256 μM). For all treatment groups, MTT cell viability (**LHS** of figure) and PI cell death (**RHS** of figure) assays were performed. * Signifies that value is significantly different than that of the vehicle control for that venom as determined by a one-way ANOVA followed by Dunnett’s multiple comparisons test (*P* < 0.05, n ≥ 3). ANOVA statistics for individual statistically analysed graphs are: (**a**) F(3,10) = 3.969, P=0.0422; (**b**) F(3,10) = 10.14, P=0.0022; (**c**) F(3,12) = 29.20, P<0.0001; (**d**) F(3,10) = 4.677, P=0.0273. Error bars represent SD of at least three independent trials, and the individual values for each trial are shown as points within each of the graphs.

### Varespladib potentiates the inhibitory activity of marimastat against *B. asper* venom when used in combination

Although the findings described in **Figs. 2 & 3** suggest that the cytotoxic activity of the viper venoms under study is primarily mediated by SVMP toxins, we wanted to determine whether PLA_2_ inhibition by varespladib could potentiate the cytoprotective properties of the SVMP-inhibiting drugs DMPS and marimastat in a representative venom abundant in PLA_2_ toxins. Thus, we repeated the MTT and PI assays using *B. asper* venom and compared the protective effects of combination treatments with those conferred by individual drug therapies. While no drug-potentiation effect was observed when varespladib was combined with DMPS (**Fig. 4a-b**), when combined with marimastat such potentiation was apparent as the potency of *B. asper* venom was significantly reduced compared to the marimastat-alone treatment, as measured with both MTT and PI assays (**Fig. 4c-d**).

**Fig. 4.**
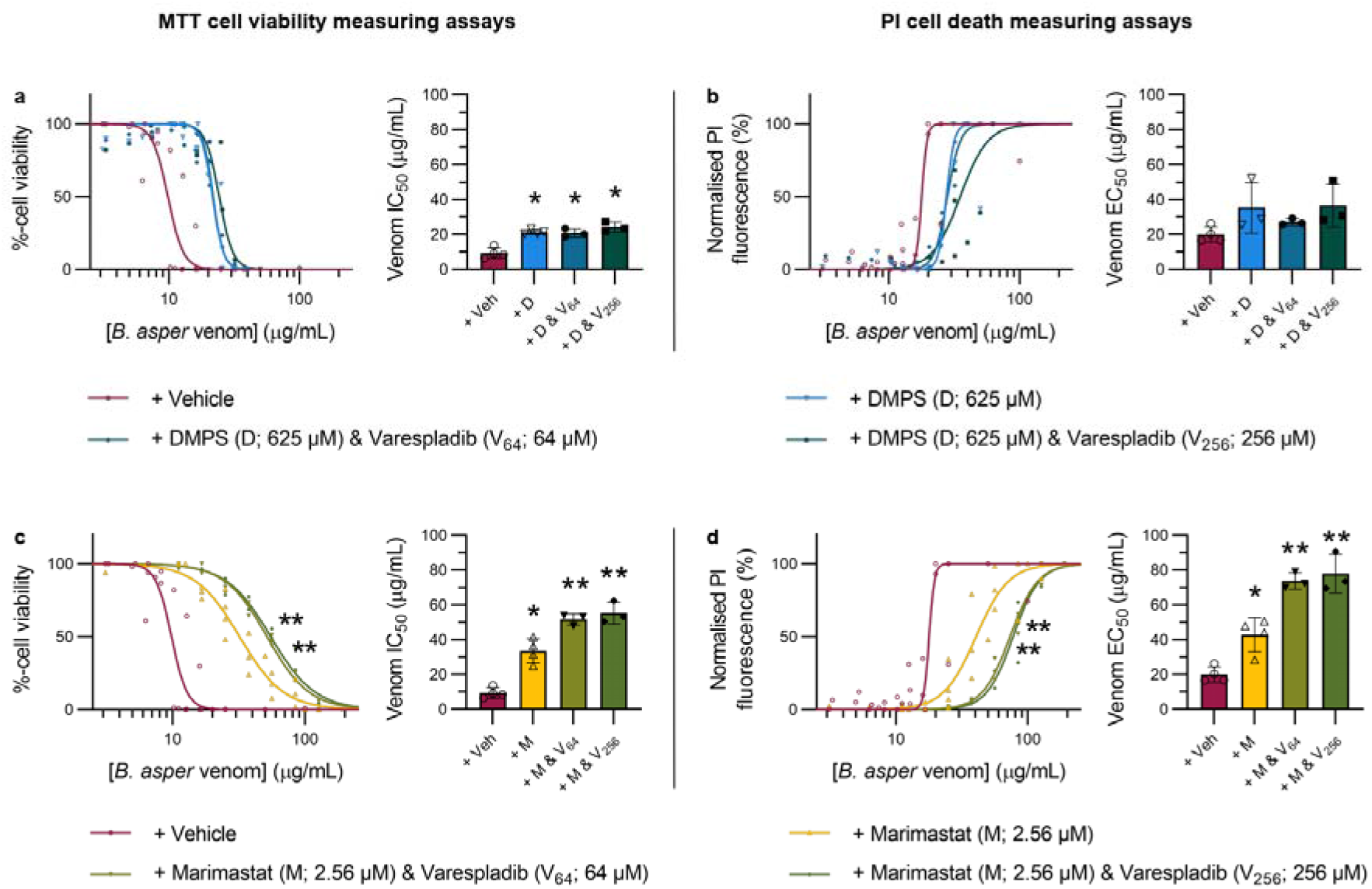
Varespladib potentiates the inhibitory effects of marimastat, but not DMPS, against *B. asper* venom in HaCaT cells. HaCaT cells were treated for 24 hours with serial dilutions of *B. asper* venom (2.2 – 190 μg/mL) that had been pre-incubated with drug vehicle control or with drug combination therapies consisting of DMPS (625 μM) plus varespladib (64 or 256 μM, abbreviated V_64_ or V_256_, respectively; **top row**) or marimastat (2.56 μM) plus V_64_ or V_256_ (**bottom row**). For all treatment groups, MTT cell viability (**LHS** of figure) and PI cell death (**RHS** of figure) assays were performed. * Signifies the value is significantly different than that of the vehicle control and ** signifies the value is significantly different than that of the marimastat-alone treatment, as determined by a one-way ANOVA comparing all treatments to each other followed by Tukey’s multiple comparisons test (*P* < 0.05, n ≥ 3). ANOVA statistics for individual statistically analysed graphs are: (**a**) F(3,10) = 26.63, P<0.0001; (**b**) F(3,9) = 2.382, P=0.1371; (**c**) F(3,10) = 56.55, P<0.0001; (**d**) F(3,10) = 40.41, P<0.0001. Error bars represent SD of at least three independent trials, and the individual values for each trial are shown as points within each of the graphs.

### Toxin-inhibiting drugs species-specifically reduce the formation of venom-induced dermal lesions *in vivo*, while drug combinations provide broad pan-species efficacy

An *in vivo* experimental animal model was used to assess the preclinical efficacy of the three toxin-inhibitory drugs and the corresponding rationally selected drug combinations at preventing the formation of venom-induced dermal lesions. We first used this model (based on the minimum necrotic dose [MND] model^53^) to determine appropriate intradermal (ID) doses of *B. asper* and *C. atrox* venom that elicit the formation of sufficiently large dermal lesions without causing any evident systemic envenoming effects, which we found to be 150 and 100 μg, respectively (**Supplementary Fig. 2**). A 39 μg dose of *E. ocellatus* venom was previously determined^54^. Next, we co-incubated the venom doses or PBS vehicle control with drug vehicle control (98.48% PBS, 1.52% DMSO), DMPS (110 μg), marimastat (60 μg), varespladib (19 μg), DMPS & varespladib (110 and 19 μg, respectively), or marimastat & varespladib (60 and 19 μg, respectively) for 30 minutes at 37 °C, prior to ID-injecting the venom-plus-drug treatments into separate groups of five mice each. After 72 hours (unless otherwise indicated) the mice were euthanised, their skin lesions excised, photographed, and measured. Representative and full image set of the resulting lesions are shown in **Fig. 5a** and **Supplementary Fig. 3**, respectively. No lesions were observed in the drug-only controls (**Fig. 5b)**. *B. asper* venom caused a mean lesion area of 41.9 mm^2^ which, in contrast to the cell data, was not significantly reduced by marimastat (55.1 mm^2^) but was by varespladib (12.2 mm^2^). Although DMPS (21.1 mm^2^) visually appeared to reduce the mean lesion area caused by *B. asper* venom, this was not statistically significant (P=0.1535) (**Fig. 5c**). *C. atrox* venom caused a mean lesion area of 19.1 mm^2^, which was significantly reduced in size by all three drug treatments: DMPS (3.1 mm^2^), marimastat (4.4 mm^2^) and, again in contrast to the cell data, varespladib (5.8 mm^2^) (**Fig. 5d**). *E. ocellatus* venom caused a mean lesion area of 5.0 mm^2^. In contrast with the other two venoms, varespladib was ineffective at reducing the lesion size (7.0 mm^2^). Both SVMP inhibitors appeared to substantially reduce *E. ocellatus* venom-induced lesions, with all five marimastat-treated and four of the five DMPS-treated mice displaying no lesions; however, only marimastat’s effects were significant (0 mm^2^) while those of DMPS were not, due to the single outlier value in this treatment group (1.0 mm^2^, P=0.0856) (**Fig. 5e**).

**Fig. 5.**
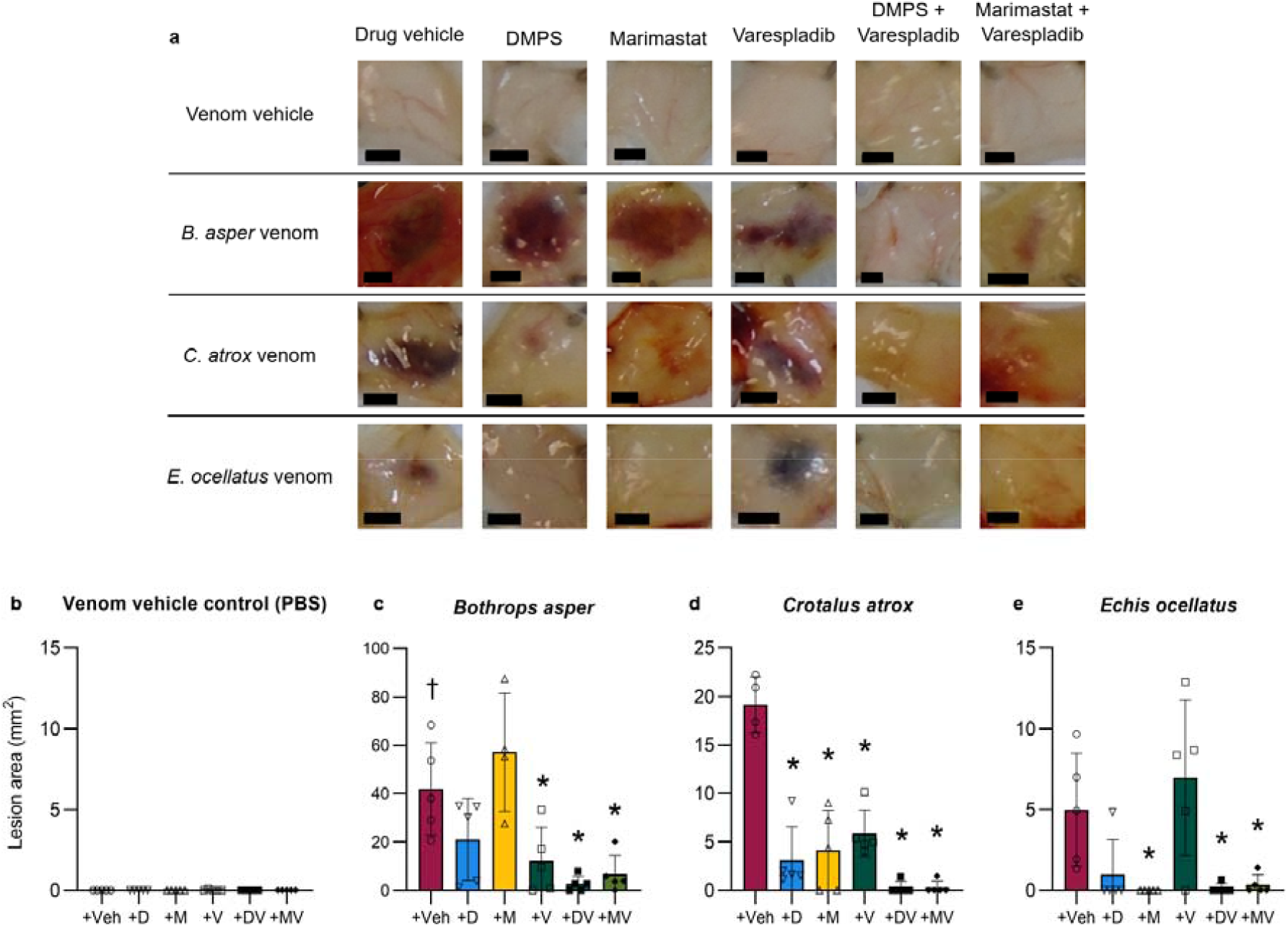
Dermal lesions induced by distinct snake venoms are inhibited by drug combinations containing an SVMP and a PLA_2_ inhibitor. Mice (n = 4-5) were ID injected with *B. asper* (150 μg), *C. atrox* (100 μg), or *E. ocellatus* (39 μg) venom or venom vehicle control (PBS) that had been pre-incubated with drug vehicle control (98.48% PBS, 1.52% DMSO; Veh), DMPS (110 μg; D), marimastat (60 μg; M), varespladib (19 μg; V), DMPS & varespladib (110 and 19 μg, respectively; DV), or marimastat & varespladib (60 and 19 μg, respectively; MV). After 72 hours^†^ the mice were euthanised and their lesions excised, height and width measured with callipers, and photographed. (**a**) Representative images of the lesions resulting from each treatment group (black scale bar represents 3 mm). Bar graphs summarising the average total lesion areas for each drug treatment group when pre-incubated with (**b**) venom vehicle control (PBS), (**c**) *B. asper*, (**d**) *C. atrox*, or (**e**) *E. ocellatus* venom. † Signifies that these mice were culled at 24 h instead of the usual 72 h, due to their external lesions progressing to the maximum permitted size defined in our animal ethics licence, thus resulting in early euthanasia. * Signifies that value is significantly different than that of the drug vehicle control for that venom as determined by a one-way ANOVA followed by Dunnett’s multiple comparisons test (*P* < 0.05, n ≥ 4). ANOVA statistics for individual statistically analysed graphs are: (**b**) F(5,24) = 1.000, P=0.4389, (**c**) F(5,23)=8.808, P<0.0001; (**d**) F(5,23) = 28.80, P<0.0001; (**e**) F(5,24) = 6.587, P=0.0005. Error bars represent SD of at least 4 lesions, and the individual values for each lesion are shown as points within each of the bars.

Using the same *in vivo* methods, we then tested combination therapies consisting of the PLA_2_-inhibiting varespladib with the SVMP-inhibiting DMPS or marimastat against these same three venoms. In contrast to the single drug therapies, which displayed variable efficacies depending on the snake species and rarely completely inhibited lesion formation in individual mice, both combination therapies significantly inhibited lesion formation caused by all three venoms tested, with many individual mice displaying no lesion development at all (**Fig. 5, Supplementary Fig. 3**). Thus, mean *B. asper* venom-induced lesions (41.9 mm^2^) were decreased to 2.7 and 6.7 mm^2^ (**Fig. 5c)**, *C. atrox* lesions (19.1 mm^2^) to 0.3 and 0.3 mm^2^ (**Fig. 5d**), and *E. ocellatus* lesions (5.0 mm^2^) to 0.1 and 0.4 mm^2^ (**Fig. 5e**) by the DMPS-plus-varespladib (DV) and marimastat-plus-varespladib (MV) combination therapies, respectively.

### Histopathological analysis of lesions confirms SVMP- and PLA_2_-inhibiting drugs and their combinations protect against snake venom-induced dermonecrosis

To better understand the dermal pathology induced by the snake venoms *in vivo* with and without co-incubation with DMPS, marimastat, varespladib, or their combinations, sections were prepared from formalin-fixed, paraffin-embedded skin lesion (or healthy control) tissue and stained with haematoxylin & eosin (H&E) dye. Photomicrographs were taken of each section at 100X magnification (10X objective, 10X ocular) for analysis and a severity scoring system was developed, which expanded upon the recent work of Ho *et al*^55^. The severity of dermonecrosis within each skin layer (epidermis, dermis, hypodermis, panniculus carnosus, and adventitia) was scored between 0 and 4 by two blinded experimenters, with 0 representing 0% of the layer within the image being affected, 1 representing up to 25%, 2 representing between 25-50%, 3 representing between 50-75%, and 4 being the most severe and representing >75% of the skin layer (**Supplementary Fig. 4**). An overall dermonecrosis score was then calculated from the mean of the resulting scores obtained for the various layers (**Fig. 6**). Representative photomicrographs of no, partial, and heavy dermonecrosis are shown in **Fig. 6a-c**.

**Fig. 6.**
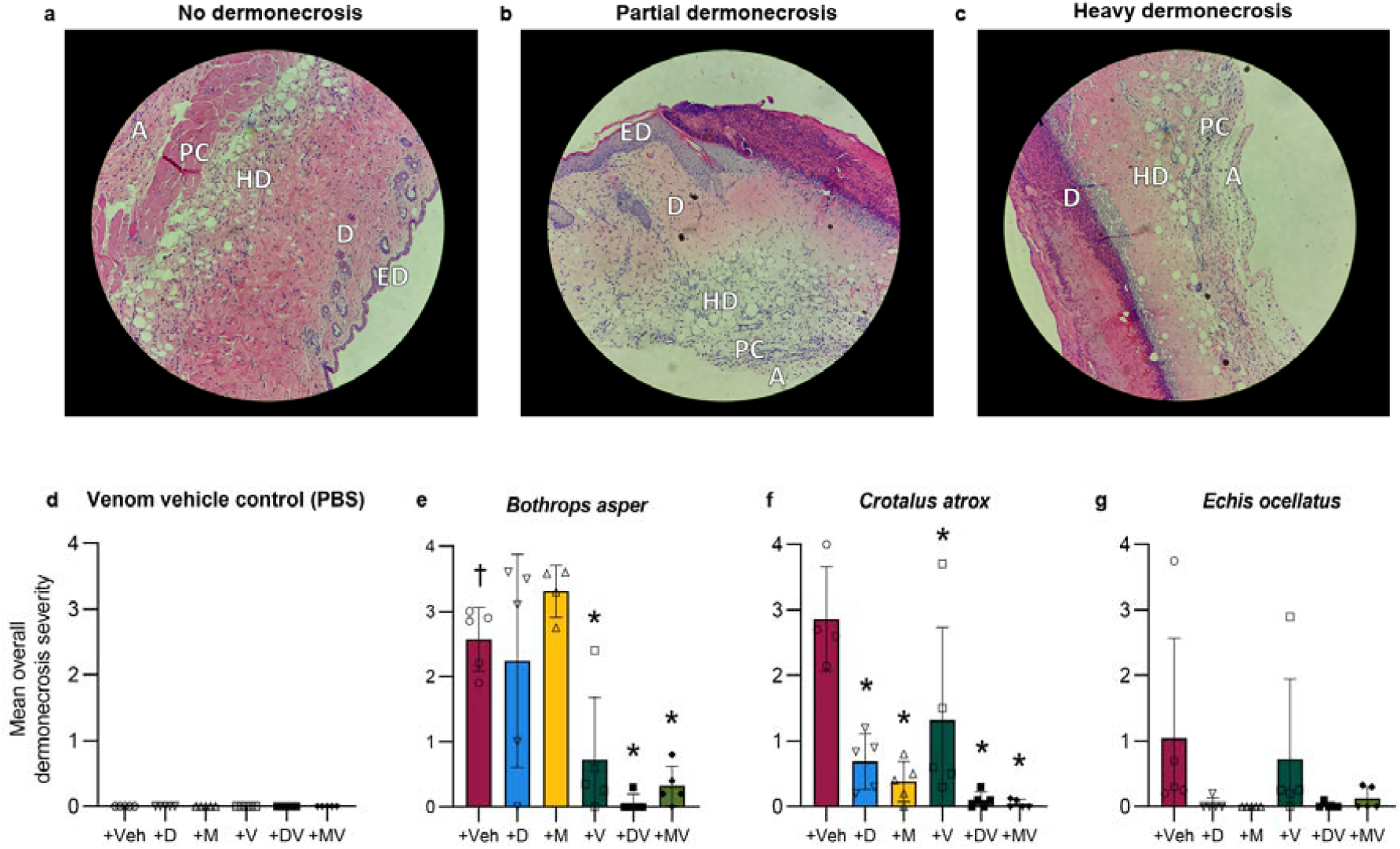
Histopathological analysis of ID-injection site cross-sections confirms venom-induced dermonecrosis can be reduced using SVMP- and PLA_2_-inhibiting drugs. Four μm H&E sections were prepared from formalin-fixed, paraffin-embedded tissue from dermal injection sites and photographed at 100X magnification. Two blinded and independent experimenters scored, between 0-4, the percentage of each skin layer that was necrotic (0=0%, 1=0-25%, 2=25-50%, 3=50-75%, and 4=75-100%). The highest recorded score per cross-section was used as a measure of the maximum severity reached within each skin sample. Representative 100X-magnified images showing (**a**) no dermonecrosis (mean overall dermonecrosis score of 0), (**b**) partial dermonecrosis (1.4) and (**c**) heavy dermonecrosis (2.4), with epidermis (ED), dermis (D), hypodermis (HD), panniculus carnosus (PC), and adventitia (A) annotated in each image (note that the ED is not visible in the ‘Heavy dermonecrosis’ image due to the severity of the ulceration, and was therefore given a necrosis score of 4). Bar graphs summarising the mean overall dermonecrosis severity scores in cross-sections from mice ID-injected with (**d**) venom vehicle control (PBS), (**e**) *B. asper* venom, (**f**) *C. atrox* venom, or (**g**) *E. ocellatus* venom that had been pre-incubated with drug vehicle control (98.48% PBS, 1.52% DMSO; Veh), DMPS (110 μg; D), marimastat (60 μg; M), varespladib (19 μg; V), DMPS-plus-varespladib (110 and 19 μg, respectively; DV), or marimastat-plus-varespladib (60 and 19 μg, respectively; MV). † Signifies these mice were culled at 24 h instead of the usual 72 h, due to their external lesions progressing to the maximum permitted size defined in the animal ethics licence, resulting in early euthanasia. * Signifies that value is significantly different than that of the drug vehicle control as determined by a one-way ANOVA followed by Dunnett’s multiple comparisons test (*P*<0.05, n≥4). ANOVA statistics: (**e**) F(5,23) = 11.81, P<0.0001; (**f**) F(5, 23)=10.30, P<0.0001; (**g**) F(5,24)1.531, P=0.2178. Error bars represent SD of at least 4 scores, and individual scores are shown as points within each of the figures’ bars.

The drug treatments plus venom vehicle control induced no dermonecrosis (**Fig. 6d and Supplementary Fig. 4a**). The varespladib, DV, and MV treatments decreased *B. asper* venom-induced dermonecrosis in the epidermis, dermis, hypodermis, and panniculus carnosus layers, though not in the adventitia, while neither DMPS nor marimastat alone inhibited the effects of *B. asper* venom in any of the skin layers (**Supplementary Fig. 4b**).

This collectively resulted in the varespladib, DV, and MV treatments decreasing the overall mean dermonecrosis score induced by *B. asper* venom from 2.57 to 0.72, 0.06, and 0.32, respectively, while DMPS and marimastat were ineffective (**Fig. 6e**). All treatments decreased *C. atrox* venom-induced dermonecrosis in the epidermis and dermis, and all but varespladib did so in the hypodermis, though no treatment had a significant effect in the panniculus carnosus or adventitia (**Supplementary Fig. 4c**). This resulted in the DMPS, marimastat, varespladib, DV, and MV treatments decreasing the overall mean dermonecrosis score induced by *C. atrox* venom from 2.86 to 0.69, 0.38, 1.32, 0.10, and 0.04, respectively (**Fig. 6f**). Lastly, the marimastat, DV, and MV treatments significantly decreased *E. ocellatus* venom-induced dermonecrosis only in the dermis while DMPS and varespladib did not; no significant results were calculated from any treatment in any other skin layer (**Supplementary Fig. 4d**). While the mean overall dermonecrosis score induced by *E. ocellatus* venom was not significantly decreased by any treatment, there was a trend towards inhibition with DMPS, marimastat, DV, and MV resulting in mean overall dermonecrosis scores of 0.04, 0.00, 0.02, and 0.12, respectively, versus 1.04 for the drug-vehicle control and 0.74 for the varespladib treatment (**Fig. 6g**). Note that minimal necrosis was observed in the adventitia even in the absence of drug treatment, suggesting that histological scoring of necrosis in this layer is likely less informative than in other skin layers.

## Discussion

Antivenom remains the only currently available specific treatment for snakebite envenoming. Despite being lifesaving therapies, antivenoms have a number of limitations that hamper their clinical utility, and thus new treatments with improved pan-snake species effectiveness, safety, and affordability are sorely needed^8,30,31^. Of particular importance is the need to develop effective treatments for tackling snakebite-induced local tissue damage, for which current antivenoms are minimally effective^1,20^. Due to their smaller size and pharmacological properties that result in superior tissue distribution versus large antibodies, small molecule drugs may offer a more effective way of preventing morbidity-causing peripheral tissue damage around the bite-site that is typical of cytotoxic snakebite envenoming^8,25,30–32^. The properties of small molecule drugs could be further exploited by developing oral snakebite therapies to be administered in the field immediately after a victim is bitten, or topical therapies that could be applied directly to the bite-site. The latter application could perhaps better ensure that the drug(s) could reach the cytotoxins in the event that venom-induced vasculature destruction prevented orally- or IV-administered drugs from reaching the tissue where the bite occurred. Both administration methods could rapidly reduce the damaging effects of a snakebite almost immediately after the bite occurs, buying the victim time to reach hospital for IV-administered antivenom to complete their treatment^8,25,26,29–32^.

In this study we sought to determine whether three toxin-inhibiting small molecule drugs (DMPS, marimastat, and varespladib), all of which have exhibited promising neutralising capabilities against snake venom-induced systemic effects previously^23,26–28,35–41^, were capable of effectively preventing snake venom-induced dermonecrosis and thus might show promise for future translation as novel treatments of local tissue damage following snakebite envenoming. Cell-based assays were selected as higher throughput and ethically acceptable alternatives to *in vivo* experiments for initial toxin-inhibitory experiments. The MTT assays were used to detect two different effects of venoms on keratinocytes in culture, i.e., cell viability-inhibition and cellular detachment. Both effects are relevant in terms of the pathology induced by venoms in the skin. First, we determined the potency of a panel of geographically diverse and taxonomically distinct medically important snake species (both viperids and elapids) in HaCaT cells^44,45^, with resulting IC_50_ values showing that the majority of the venoms (9 of the 11 tested) were equipotently cytotoxic (**Fig. 1)**. These findings were unexpected given the extensive variation in toxin composition among these snake species^9,56^. While reductions in cytotoxic potency observed with *D. russelii* were minor, *N. haje* venom was significantly less cytotoxic than all other venoms, including those from the congeneric spitting cobra species *N. nigricollis* and *N. pallida;* however, this observation is in line with the distinct, predominantly neurotoxic, composition and functional activity of this non-spitting cobra venom^56,57^. As an additional pharmacological measure, the Hill slopes of all venoms were calculated and compared (**Fig. 1m**), and while no slope was significantly different from another, the magnitudes of all 11 were greater than 1.5 and thus considered ‘steep’^58^, meaning a small change in venom concentration can lead to a large change in overall pathological effect. In classical pharmacology, steep Hill slope values can be explained by the activity of one bioactive factor (e.g. a toxin) agonising the activity of another, thus allowing them to reach maximum pathological efficacy over a shorter concentration range. This indicates a phenomenon known as ‘positive cooperativity’^47,48^ which suggests probable pathological synergy between certain snake venom toxins, something that has been previously evidenced^59–61^.

Our skin cell assays demonstrated that the SVMP-inhibitors DMPS and marimastat may be effective anti-cytotoxic drugs as individual therapies, although their inhibitory effects were not universal across all cytotoxic snake venoms (**Fig. 2**). Unexpectedly, the PLA_2_ inhibitor varespladib was ineffective against any of the venoms tested, despite it displaying impressive results against systemic venom-induced toxicity previously^23,28,38–41^. To explore whether MTT assays are simply a poor assay choice for testing PLA_2_-inhibitors against cytotoxic venoms, we multiplexed them with a secondary cytotoxicity assay using PI to measure cell membrane disruption^51,52^. Nevertheless, varespladib remained ineffective in these assays, suggesting that much of the cytotoxicity observed in these studies is mediated by SVMP toxins rather than PLA_2_s (**Fig. 2** and **3**); however, when we treated the cells with varespladib in combination with marimastat we observed significant reductions in the potency of *B. asper* venom versus the marimastat-alone treatment (**Fig. 4**). These findings suggest that PLA_2_ toxins may indeed, at least to some extent, contribute to cytotoxic venom effects, and that combining an SVMP-inhibitor with a PLA_2_-inhibitor may improve overall treatment efficacy. Interestingly, this anti-cytotoxic potentiation of marimastat by varespladib was not observed with DMPS despite this drug also being a SVMP-inhibitor. This dichotomy is likely due to the fact that these drugs’ mechanisms of action are different, as marimastat directly inhibits metalloproteinases by acting as a peptidomimetic and binding covalently to the Zn^2+^ ion present in the active site^25,37,62–64^, while the inhibitory mechanism of action of DMPS is solely the result of Zn^2+^ chelation^26,62^. These mechanistic variations likely underpin the previously described differences in SVMP-inhibiting potencies of these drugs *in vitro*^27,65,66^.

For subsequent *in vivo* neutralisation experiments, we tested three venoms whose cytotoxic potencies were reduced by both DMPS and marimastat in the cell assays, that were sourced from three different genera of snakes displaying considerable inter-species toxin variability^9^, and that inhabit distinct localities: *B. asper* (Latin America), *C. atrox* (North America), and *E. ocellatus* (West Africa). Using a drug pre-incubation^26,27^ model of venom dermonecrosis in mice^53^, we showed that DMPS, marimastat, and varespladib are each individually capable of inhibiting venom-induced skin lesion formation *in vivo*, though the efficacy of each drug was restricted to only certain venoms (**Fig. 5**). Thus, in line with the cell cytotoxicity findings, DMPS significantly reduced dermonecrotic lesions induced by *C. atrox* venom, while marimastat was effective against both *C. atrox* and *E. ocellatus*. However, contrasting with our cell data, at the therapeutic doses tested marimastat did not significantly reduce lesions caused by *B. asper* venom, nor did DMPS against *B. asper* or *E. ocellatus* venom. Further, varespladib was effective at significantly inhibiting *B. asper* and *C. atrox* venom *in vivo*, though not *E. ocellatus* venom, likely due to this venom being dominated by SVMPs^9^. In summary these findings clearly evidence that, while they are undoubtedly informative and could be valuable screening tests, cell-based cytotoxicity assays do not fully recapitulate findings obtained through *in vivo* dermonecrosis experiments, and thus additional models are required downstream to robustly assess drug efficacy against this aspect of local snakebite envenoming.

While the results of the single drug trials against select cytotoxic snake venoms *in vivo* are certainly promising, they also evidence how single drugs are not paraspecifically effective. In contrast, the two combination therapies tested (MV and DV) both effectively and substantially reduced the development of macroscopic dermal lesions caused by all three tested venoms despite major differences in their toxin compositions^9^, with many of the drug combination-treated mice displaying no dermal lesion development at all (**Fig. 5, Supplementary Fig. 3**). This could be explained by the variable roles that SVMPs and PLA_2_s play in the pathogenesis of skin damage induced by different snake venoms.

Histopathological analysis of the resulting lesions confirmed the efficacy of the drug combinations with both DV and MV significantly reducing the severity of overall dermonecrosis observed in the skin layers of mice envenomed with *B. asper* and *C. atrox* venoms, albeit the results in mice injected with *E. ocellatus* venom were not significant due to the dose being too low to consistently induce sufficient dermonecrosis in the drug-vehicle control group (**Fig. 6**). These data strongly evidence the utility of small molecule drug combination therapies targeting SVMP and PLA_2_ toxins as largely preclinically efficacious and pan-species effective therapies for the treatment of the local skin-damaging effects of viperid snakebite envenoming.

When combined with the results of Albulescu, *et al*.^27^, our findings show that combination drug therapies simultaneously targeting SVMP and PLA_2_ toxins are likely to be useful for tackling both the life-threatening systemic and morbidity-causing local pathologies caused by diverse viperid snake venoms. Because snakebite is a global health challenge that predominately affects populations in lower- and middle-income countries (LMICs), our findings here have considerable consequences for the future treatment of this WHO priority-listed NTD, particularly when considering that *E. ocellatus* are responsible for most snakebite deaths in West Africa^67^ and *B. asper* causes the vast majority of severe snakebites in Central America^68^. Further, evidence of inhibitory potential against *C. atrox*, a North American pit viper species, may enable a strategy for the future global translation of drug combination therapies by leveraging one of the few financially viable markets available for snakebite. Such an approach must, however, ensure that a robust access plan for LMIC communities is developed in parallel to avoid potential future distribution pitfalls, like those recently reported around the inequitable distribution of COVID vaccines^69,70^.

There remains much work to be done to translate these drugs and their combinations into approved snakebite therapies. This includes additional preclinical research, for example against the venoms of additional snake species (e.g. other viperids and cytotoxic *Naja* spitting cobras^56,71,72^), assessment in ‘challenge-then-treat’ models of envenoming^26^, trials testing different routes of therapeutic administration^26^, and experiments to better understand their pharmacokinetics and pharmacodynamics to elucidate informed dosing regimens and potential drug-drug interactions. Since a major anticipated benefit of drug therapies for snakebite is their potential to be orally or topically formulated^8^ (i.e. in contrast with intravenously-injected antivenom), considerable research effort should focus around this space to pursue the translation of safe, affordable, community-level interventions to reduce existing treatment delays in rural tropical communities, thus improving patient outcomes. To that end it is worth noting that DMPS is already undergoing Phase I clinical trials to determine both its safety and a PK-informed oral dosing regimen for snakebite indication^73^, while methyl varespladib has recently entered Phase II trials to assess its safety, tolerability, and efficacy in snakebite victims (https://clinicaltrials.gov/ct2/show/NCT04996264). These studies emphasise the growing confidence the research community has in specific small molecule drugs as novel oral treatments for snakebite envenoming, though the data presented here highlight that additional research to develop these (among other) drugs into combination therapies is likely to yield treatments with superior pan-snake species effectiveness than any single drug alone.

In conclusion, our data provide strong evidence that the small molecule drugs DMPS, marimastat, and varespladib can significantly protect against dermonecrosis associated with local snakebite envenoming, though their efficacy is limited to certain snake species. This limitation is largely overcome when the SVMP-inhibiting drugs DMPS or marimastat are used in combination with the PLA_2_-inhibiting drug varespladib, most likely due to the dual role of SVMPs and PLA_2_s in the pathogenesis of tissue damage across snake species. Our data illustrate that toxin-inhibiting small molecule drugs show considerable potential as novel broad-spectrum treatments against the local skin-damaging effects of cytotoxic snake venoms, an important outcome when considering current antivenom therapies have limited efficacy against severe local envenoming. Our findings therefore advocate for further research to translate these drugs and their combinations into community-deliverable snakebite treatments with the goal of significantly reducing the morbidity associated with one of the world’s most neglected tropical diseases.

## Methods

### Chemicals, Drugs and Biological Materials

Thiazolyl blue methyltetrazolium bromide (MTT; M5655), dimethyl sulfoxide (DMSO; 276855), and propidium iodide (PI; P4170) were purchased from Sigma-Aldrich (Merck). Dulbecco’s modified Eagle’s medium (DMEM; 11574516), foetal bovine serum (FBS; 11573397), FluoroBrite DMEM (A1896701), glutaMAX supplement (35050038), penicillin-streptomycin (11528876), phosphate buffered saline (11503387), and TrypLE Express were purchased from Gibco (Thermo Fisher Scientific). Marimastat (M2699) and varespladib (SML1100) were purchased from Sigma-Aldrich (Merck), and 2,3-dimercapto-1-propanesulfonic acid sodium salt monohydrate (DMPS; H56578) was purchased from Alfa Aesar. Working stocks were: DMPS (PBS, 400 mM, made fresh with each use from lyophilised powder), marimastat (10 mM, ddH_2_O), and varespladib (65.7 mM, DMSO).

### Venoms

Venoms were sourced from either wild-caught snakes maintained, or historical venom samples stored, in the herpetarium of the Centre for Snakebite Research & Interventions at the Liverpool School of Tropical Medicine (LSTM). This facility and its protocols for the husbandry of snakes are approved and inspected by the UK Home Office and the LSTM and University of Liverpool Animal Welfare and Ethical Review Boards. The venom pools were from snakes with diverse geographic localities, namely: *Bitis arietans* (Nigeria), *Bothrops asper* (Costa Rica [Caribbean region]), *Crotalus atrox* (captive bred [USA lineage]), *Calloselasma rhodostoma* (Malaysia), *Daboia russelii* (Sri Lanka), *Echis carinatus* (India), *Echis ocellatus* (Nigeria), *Naja haje* (Uganda), East-African *Naja nigricollis* (Tanzania), West-African *Naja nigricollis* (Nigeria), and *Naja pallida* (Tanzania). Note that the Indian *E. carinatus* venom was collected from a specimen that was inadvertently imported to the UK via a boat shipment of stone, and then rehoused at LSTM on the request of the UK Royal Society for the Prevention of Cruelty to Animals (RSPCA). Crude venoms were lyophilized and stored at 4 °C to ensure long-term stability. Prior to use, venoms were resuspended to 10 mg/ml in PBS and then kept at -80 °C until used in the described experiments, with freeze-thaw cycles kept to a minimum to prevent degradation.

### Cells

The immortalised human epidermal keratinocyte line, HaCaT^44,45^, was purchased from Caltag Medsystems (Buckingham, UK). Cells were cultured in phenol red-containing DMEM with GlutaMAX supplemented with 10% FBS, 100 IU/mL penicillin, 250 μg/mL streptomycin, and 2 mM sodium pyruvate (standard medium; Gibco) per Caltag’s HaCaT protocol. For assays that contained the fluorescent dye, PI, a medium specifically formulated for fluorescence-based cell assays was used instead: FluoroBrite DMEM supplemented with 1% GlutaMAX 100x supplement, 1% FBS, 100 IU/mL penicillin, 250 μg/mL streptomycin, and 2 mM sodium pyruvate (minimally fluorescent medium; Gibco). The cells were split and growth medium changed 2x per week up to a maximum of 30 passages. Cells were maintained in a humidified, 95% air/5% CO_2_ atmosphere at 37 °C (standard conditions).

### MTT Cell Viability and PI Cell Death Assays

MTT assays were used to evaluate the cell (HaCaT) viability-inhibiting activity of snake venoms and high concentrations of drug inhibitors and were based on the methods of Issa, *et al*.^74^. PI assays were used to evaluate the cell death and were based on the methods of Chitolie & Toescu^52^.

#### MTT assays alon

HaCaT cells were seeded (5,000 cells/well, clear-sided 96-well plates) in standard medium, then left to adhere overnight at standard conditions. The next day, serial dilutions were prepared in standard medium of (a) venom treatments (1-1,024 μg/mL; i.e. **Fig. 1**), (b) DMPS (9.8-10,000 μM), marimastat (0.04-40.96 μM), or varespladib (1-1,024 μM) treatments (i.e. **Supplementary Fig. 1**), or (c) venoms (2.5-240 μg/mL) preincubated with a single concentration (the MTC_½_ as determined in b) of DMPS (625 μM), marimastat (2.56 μM), varespladib (128 μM) or drug vehicle control at standard conditions for 30 minutes (i.e. **Fig. 2**). Cells were treated with each prepared solution (100 μL/well, triplicate wells/prepared solution) for 24 hours. Thereafter, MTT solution (3.33 mg/mL) was prepared in PBS, filtered through a 0.22 μm syringe filter, then 30 μL added to each treatment well (and to ‘no treatment’ positive control wells and ‘no cell’ negative control wells) creating a final MTT concentration of 0.833 mg/mL. The plates were then incubated for 1.5 h at standard conditions for the MTT reaction to occur, after which medium was aspirated from all wells and replaced with 100 μL of DMSO. Plates were shaken to ensure a homogenous mixture of purple formazan, and then absorbance (550 nm; A_550_) read on a FLUOstar Omega Microplate Reader. The % adherent cell viability for each treatment well was calculated as follows:

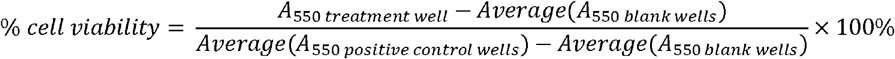

The concentration that resulted in a 50% reduction in adherent cell viability (IC_50_) was calculated from the log_10_ concentration versus normalised response curves using the ‘log(inhibitor) vs. normalized response – Variable slope’ in GraphPad Prism, which uses the following equation:

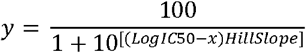

where *y* is the normalised %-cell viability values and *x* is the log_10_ of the venom concentrations.

#### MTT assays multiplexed with PI assays

HaCaT cells were seeded (20,000 cells/well, black-sided & clear-bottomed 96-well plates) in standard medium, then left to adhere overnight at standard conditions. The next day, serial dilutions of *D. russelii* or *B. asper* venom (2.2-127 μg/mL) with a single concentration of DMPS (625 μM), marimastat (2.56 μM), varespladib (256 μM), DMPS & varespladib (DV; 625 μM and 256 μM, respectively), marimastat & varespladib (MV; 2.56 μM and 256 μM, respectively) or drug vehicle control (i.e. **Fig. 3 & 4**) were prepared in minimally fluorescent medium supplemented with 74.8 μM (50 μg/mL) PI and pre-incubated at standard conditions for 30 minutes prior to cell exposure. After pre-incubation, cells were treated with each prepared solution (100 μL/well, triplicate wells/prepared solution). After 24 h, PI fluorescence (Ex_544_/Em_612_, read from bottom of plate at multiple points within each well) was read on a FlexStation 3 Multi-Mode Microplate Reader (Molecular Devices). PI relative fluorescence units (RFUs) of each treatment minus those of the PI solution blanks (no cells) were recorded as a measure of cell death and normalised between 0-100 to create PI dose-response curves. The venom dose at which the normalised PI reading was 50% of each treatment’s maximum (the half maximal effective concentration, or EC_50_ value) was determined using the ‘log(agonist) vs. normalized response – Variable slope’ in GraphPad Prism, which uses the following equation:

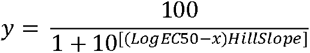

where *y* is the normalised PI (RFU_treatment_ minus RFU_blanks_) values and *x* is the log_10_ of the venom concentrations. After the PI assays were completed, the PI-containing treatment solutions were aspirated from each well and replaced with 100 μL/well of minimally fluorescent medium containing 0.833 mg/mL of filtered MTT solution, and MTT assays completed and analysed as described above.

### Animal ethics and maintenance

All animal experiments were conducted using protocols approved by the Animal Welfare and Ethical Review Boards of the Liverpool School of Tropical Medicine and the University of Liverpool and were performed in pathogen-free conditions under licensed approval (PPL #P58464F90) of the UK Home Office and in accordance with the Animal [Scientific Procedures] Act 1986 and institutional guidance on animal care. All experimental animals (18-20 g [4-5 weeks old], male, CD-1 mice, Charles River, UK) were acclimatised for a minimum of one week before experimentation with their health monitored daily. Mice were grouped in cages of five, with room conditions of approximately 22 °C at 40-50% humidity, with 12/12 hour light cycles, and given *ad lib* access to CRM irradiated food (Special Diet Services, UK) and reverse osmosis water in an automatic water system. Mice were housed in specific-pathogen free facilities in Techniplast GM500 cages containing Lignocell bedding (JRS, Germany), Sizzlenest zigzag fibres as nesting material (RAJA), and supplied with environmental enrichment materials.

### Preclinical anti-dermonecrosis efficacy of small molecule drugs *via* a preincubation model of envenoming

The *in vivo* experimental design was based upon 3R-refined WHO-recommended envenomation protocols^26,53^, with animals randomised and observers being blinded to the drug and vehicle treatments, and the anti-dermonecrosis methods were based on the Minimum Necrotizing Dose (MND) principles originally described in Theakston and Reid^53^. Before commencing the drug trials, appropriate necrotic doses of *B. asper* and *C. atrox* venom-alone were determined. Groups of two-to-three mice received ID injections in the shaved rear quadrant on the dorsal side of their flank skin with 50 μL treatments containing *B. asper* (50, 75, 100, 150, 200, or 250 μg) or *C. atrox* (30.5, 50, 75, 100, 150, or 200 μg) venom. The most appropriate experimental doses were those that consistently induced visible external lesions that grew to no more than 10 mm in diameter without inducing signs of systemic envenoming, as stipulated by our ethics licencing; these were determined to be 150 μg of *B. asper* venom and 100 μg of *C. atrox* venom (i.e. **Supplementary Fig. 2**). The 39 μg dose for *E. ocellatus* venom was previously published^54^. For anti-dermonecrosis small molecule drug trials, groups of five mice received experimental doses per mouse that consisted of: (a) venom alone, (b) venom plus drug (DMPS, marimastat, varespladib, DMPS plus varespladib [DV], or marimastat plus varespladib [MV]), or (c) venom vehicle (PBS) plus drug. Albulescu, *et al*. previously used 60 μg/mouse of marimastat in their preclinical ID haemotoxicity trials^27^, therefore this same marimastat dose was chosen for our dermonecrosis trials. A slightly higher dose of DMPS (110 μg/mouse) was chosen due to our findings that DMPS is a less potent inhibitor of cytotoxicity than marimastat in HaCaT cells, and a lower dose of varespladib (19 μg/mouse) was chosen due to solubility issues at higher drug concentrations. Stock solutions of DMPS and marimastat were dissolved in PBS, while the more hydrophobic varespladib was dissolved in DMSO; therefore, for the sake of inter-treatment consistency the same drug-vehicle control was used within all treatments described above, which resulted in a final treatment vehicle solution of 1.52% DMSO and 98.48% PBS. All experimental doses were prepared to a volume of 50 μL and incubated at 37 °C for 30 minutes, then kept on ice for no more than 3 hours until the mice were injected. For dose delivery, mice were briefly anesthetised using inhalational isoflurane (4% for induction of anaesthesia, 1.5-2% for maintenance) and ID-injected in the shaved rear quadrant on the dorsal side of the flank skin with the 50 μL treatments. The mice were observed three times daily up to 72 hours post-injection to check for symptoms of systemic envenomation or excessive external lesion development, which would have necessitated early termination of the animal due to reaching a humane endpoint defined by the animal ethics licence. At the end of the experiments (72 hours, except for the single group of *B. asper* venom plus drug vehicle control-treated mice that experienced greater-than-anticipated lesion development for which the time point was 24 hours), the mice were euthanised using rising concentrations of CO_2_, after which the skin surrounding the injection site was dissected and internal skin lesions measured with callipers and photographed. Cross-sections of the skin lesions were further dissected and preserved in formalin for mounting on microscopy slides for downstream histopathological analysis.

### Preparation and histopathological analysis of H&E-stained sections of venom-induced lesions

Skin samples underwent tissue processing using a Tissue-Tek VIP (vacuum infiltration processor) overnight before being embedded in paraffin (Ultraplast premium embedding medium, Solmedia, WAX060). Next, 4 μm paraffin sections were cut on a Leica RM2125 RT microtome, floated on a water bath and placed on colour slides (Solmedia, MSS54511YW) or poly-lysine slides (Solmedia MSS61012S) to dry. For haematoxylin & eosin (H&E) staining, slides were dewaxed in xylene and rehydrated through descending grades of ethanol (100%, 96%, 85%, 70%) to distilled water before being stained in haematoxylin for 5 mins, “blued” in tap water for 5 mins, then stained in eosin for 2 mins. Slides were then dehydrated through 96% and 100% ethanol to xylene and cover slipped using DPX (Cellpath SEA-1304-00A). Haematoxylin (Atom Scientific, RRBD61-X) and Eosin (TCS, HS250) solutions were made up in house. Brightfield images of the H&E-stained lesions were taken with an Echo Revolve microscope (Settings: 100x magnification; LED: 100%; Brightness: 30; Contrast: 50; Colour balance: 50), with at least five images taken per cross-section. Histologic evidence of necrosis was assessed separately for the epidermis, dermis, hypodermis, panniculus carnosus, and adventitia. Features of necrosis included loss of nuclei, nuclear fragmentation (karyorrhexis), nuclear shrinkage and hyperchromasia (pyknosis), loss of cytoplasmic detail with hypereosinophilia, loss of cell borders, and, in the case of severe necrosis, disarray with complete loss of architecture and hyalinization. In the epidermis, ulceration with superficial debris was interpreted as evidence of necrosis. In the dermis, loss of skin adnexal structures (e.g. hair follicles and sebaceous glands) and extracellular matrix disarray were also interpreted as evidence of necrosis. Expanding upon methods originally published by Ho, *et al*.^55^, the %-necrosis of each skin layer (epidermis, dermis, hypodermis, panniculus carnosus, and adventitia) within each image was assessed by two independent and blinded pathologists and scored between a 0 and 4, with a 0 meaning no observable necrosis in the layer within that image, a 1 meaning up to 25% of the layer in that image exhibiting signs of necrosis, a 2 meaning 25-50% necrosis, a 3 meaning 50-75%, and a 4 meaning more than 75% exhibiting indicators of necrosis. The mean scores of the pathologists for each layer from each image were determined, and the highest scores-per-mouse used for our data analysis as these represented the maximum necrotic severity within each lesion (i.e. **Supplementary Fig. 4**). The ‘mean overall dermonecrosis severity’ was determined for each lesion by taking the mean of the individual layer scores (i.e. **Fig. 6d-g**).

### Statistical Analysis

All data are presented as mean ± standard deviation^75^ of at least three independent experimental replicates. For cell experiments, ‘n’ is defined as an independent experiment completed at a separate time from other ‘n’s within that group of experiments; all drug and/or venom treatments within an ‘n’ were completed in triplicate wells and the mean taken as the final value for that one trial. For *in vivo* experiments, ‘n’ is defined as the number of mice in that specific treatment group^76^. Two-tailed t-tests were performed for dual comparisons, one-way analysis of variances (ANOVAs) performed for multiple comparisons with one independent variable followed by Dunnett’s or Tukey’s multiple comparisons tests when the trial data were compared to a single control group or to all other groups, respectively, as recommended by GraphPad Prism, and two-way ANOVAs performed for multiple comparisons with two independent variables followed by Dunnett’s multiple comparisons tests. A difference was considered significant if *P* ≤ 0.05.

## Acknowledgements

We would like to give our thanks to Paul Rowley for maintaining the snakes at the LSTM’s herpetarium and for routine venom extractions, Dr. Cassandra Modahl and Dr. Amy Marriott for their help with animal welfare observations during the *in vivo* experimentation, Valerie Tilston and her team at the University of Liverpool for preparing the histopathology slides, and all the staff at the University of Liverpool’s Biomedical Services Unit (BSU). Funding was provided by a (i) Newton International Fellowship (NIF\R1\192161) from the Royal Society to SRH, (ii) a Sir Henry Dale Fellowship (200517/Z/16/Z) jointly funded by the Wellcome Trust and the Royal Society to NRC, (iii) a Wellcome Trust funded project grant (221712/Z/20/Z) to NRC and JK and (iv) a UK Medical Research Council research grant (MR/S00016X/1) to NRC. This research was funded in whole, or in part, by the Wellcome Trust. For the purpose of open access, the authors have applied a CC BY public copyright licence to any Author Accepted Manuscript version arising from this submission.

## Author contributions

Conceptualisation: SRH, LOA, JK, NRC

Methodology: SRH, SAR, JMG, EC, CAD, NRC

Investigation: SRH, SAR, JMG, EC, CAD, KEB, LOA, APW, NRC

Data curation: SRH

Formal analysis: SRH, SAR, JMG, NRC

Original draft preparation: SRH, NRC

Editing: All authors

## Competing interests

The authors declare no competing interests.

## Materials & Correspondence

Correspondence and requests for materials should be addressed to NRC.

## Data availability statement

The data used to create the figures and supplementary figures displayed within can be found in the Supplementary Data File. All images of the H&E-stained dermal cross-sections of the animals’ injection sites used for histopathological analysis can be downloaded at https://doi.org/10.6084/m9.figshare.19706761.v1.

## Notes

https://doi.org/10.6084/m9.figshare.19706761.v1

### Competing Interest Statement

The authors have declared no competing interest.

