## Supplementary Fig. 1 for "Repurposed drugs and their combinations prevent morbidity-inducing dermonecrosis caused by diverse cytotoxic snake venoms"

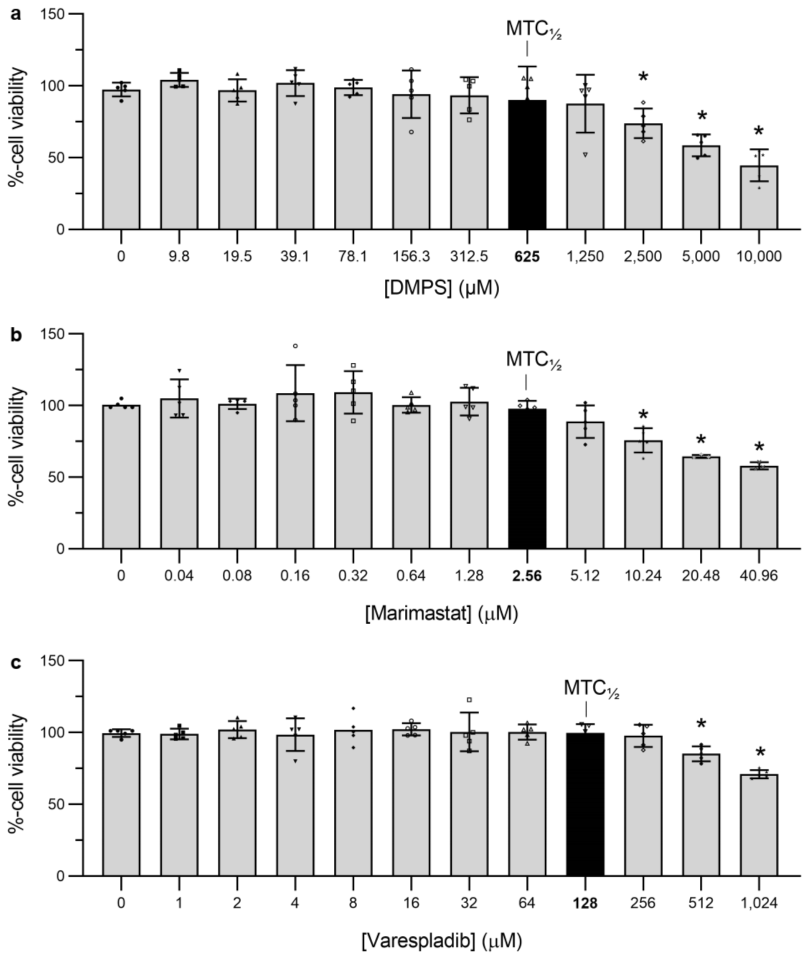


**Supplementary Fig. 1 Cell viability experiments to determine the MTC of the toxin inhibiting drugs DMPS, marimastat, and varespladib.** HaCaT cells were treated with serial dilutions of (**a**) DMPS (9.8 – 10,000 μM), (**b**) marimastat (0.04 – 40.96 μM), or (**c**) varespladib (1 – 1,024 μM) for 24 hours after which MTT assays were completed as a measure of cell viability. The maximum tolerated concentration (MTC) was defined as the highest concentration of drug that did not significantly reduce cell viability compared to the vehicle control (0), and one-half of this concentration (MTC½) was then used for all subsequent assays using these drugs unless otherwise indicated. * Signifies value is significantly lower than that of the vehicle control within each figure as determined by a one-way ANOVA followed by Dunnett's multiple comparisons test ($P < 0.05$, n ≥3). ANOVA statistics for individual graphs are: (**a**) $F(11,48) = 10.88$, $P<0.0001$; (**b**) $F(11,44) = 10.59$, $P<0.0001$; (**c**) $F(11,48) = 7.863$, $P<0.0001$. Error bars represent SD of at least three independent trials, and the individual cell viability percentages for each trial are shown as points within each bar of the above graphs.

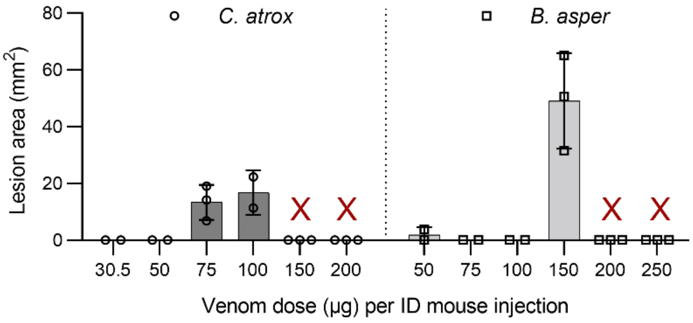


**Supplementary Fig. 2 Dermal lesion sizes determined *in vivo* in venom dose optimisation experiments.** Mice (n= 2 or 3) were ID injected with *C. atrox* (30.5, 50, 75, 100, 150 or 200 µg) or *B. asper* (50, 75, 100, 150, 200, or 250 µg) venom dissolved in PBS. After 72 hours the mice were euthanised and their lesions excised and measured with callipers. The red 'X' indicates that trialled dose was too high and that the mice had to be euthanised prior to the cessation of the experiment due to observations of systemic envenoming or excessive lesion development. Each point on each bar represents a single experimental animal, and error bars represent SD.

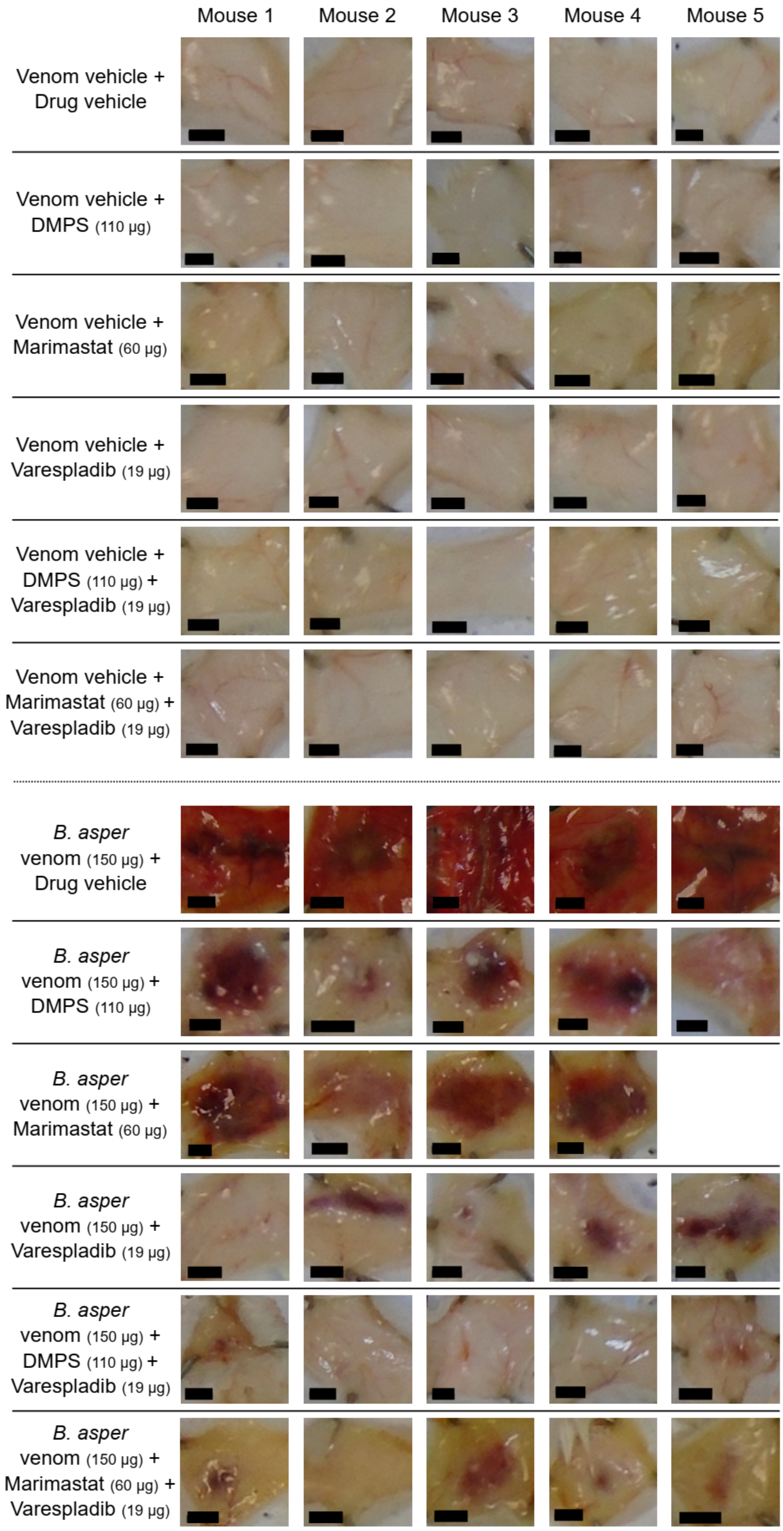

Mouse 1
Mouse 2
Mouse 3
Mouse 4
Mouse 5
Venom vehicle + Drug vehicle
Venom vehicle + DMPS (110 µg)
Venom vehicle + Marimastat (60 µg)
Venom vehicle + Varespladib (19 µg)
Venom vehicle + DMPS (110 µg) + Varespladib (19 µg)
Venom vehicle + Marimastat (60 µg) + Varespladib (19 µg)
B. asper venom (150 µg) + Drug vehicle
B. asper venom (150 µg) + DMPS (110 µg)
B. asper venom (150 µg) + Marimastat (60 µg)
B. asper venom (150 µg) + Varespladib (19 µg)
B. asper venom (150 µg) + DMPS (110 µg) + Varespladib (19 µg)
B. asper venom (150 µg) + Marimastat (60 µg) + Varespladib (19 µg)

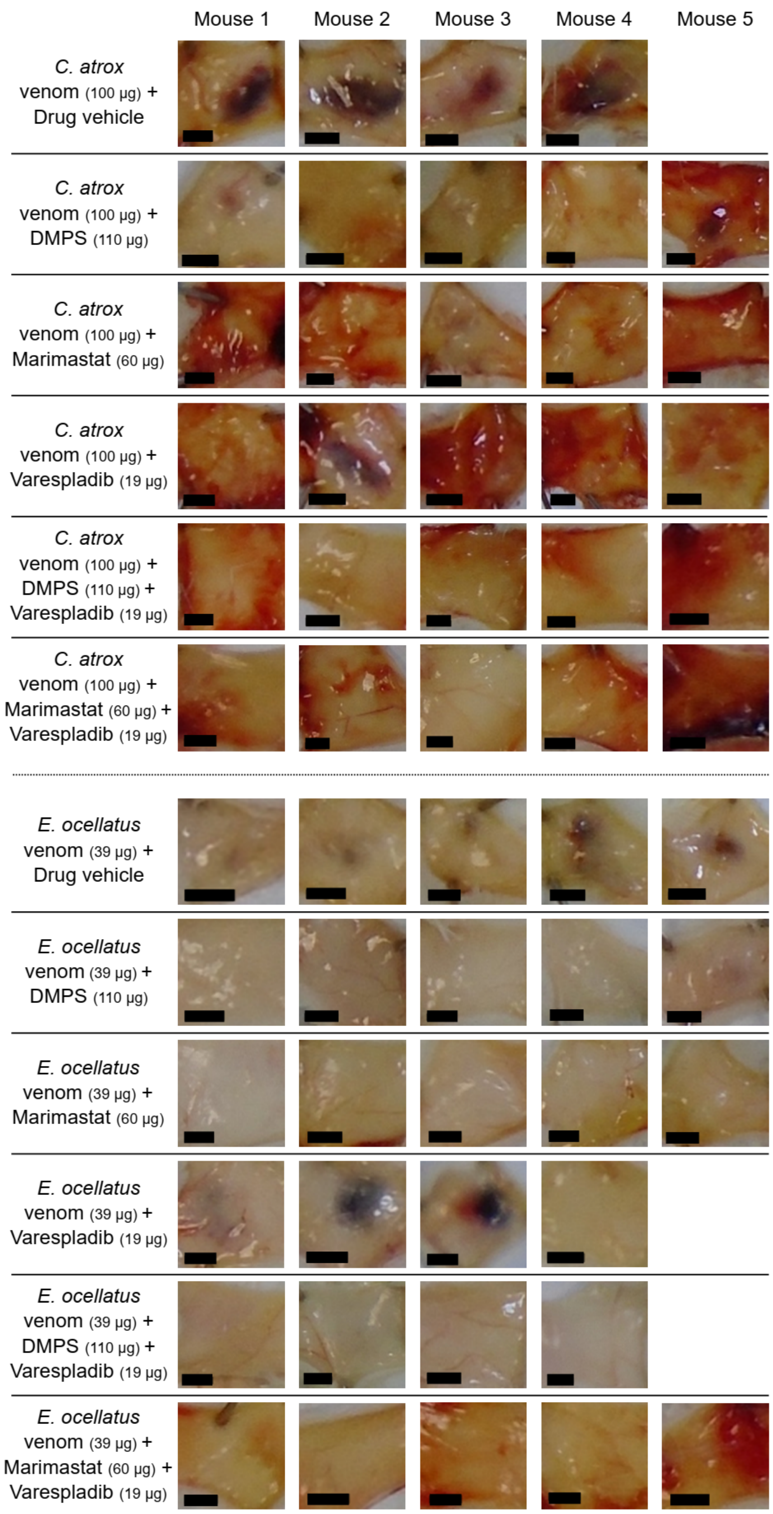


**Supplementary Fig. 3 All lesion images from every mouse in this trial, minus those that were culled before the desired timepoints due to humane endpoints being reached**. Mice were ID injected in the shaved rear quadrant on the dorsal side of the flank skin with *B. asper* (150 µg), *C. atrox* (100 µg), or *E. ocellatus* (39 µg) venom or venom vehicle control (PBS) that had been pre-incubated with drug vehicle control (98.48% PBS, 1.52% DMSO), DMPS (110 µg), marimastat (60 µg), varespladib (19 µg), DMPS & varespladib (110 and 19 µg, respectively), or marimastat & varespladib (60 and 19 µg, respectively). After 72 hours (except for the *B. asper* venom plus drug vehicle control treated mice, which were culled at 24 hours due to reaching a humane endpoint as defined in animal ethics licence) the mice were euthanised and their lesions excised, measured with callipers, and photographed.

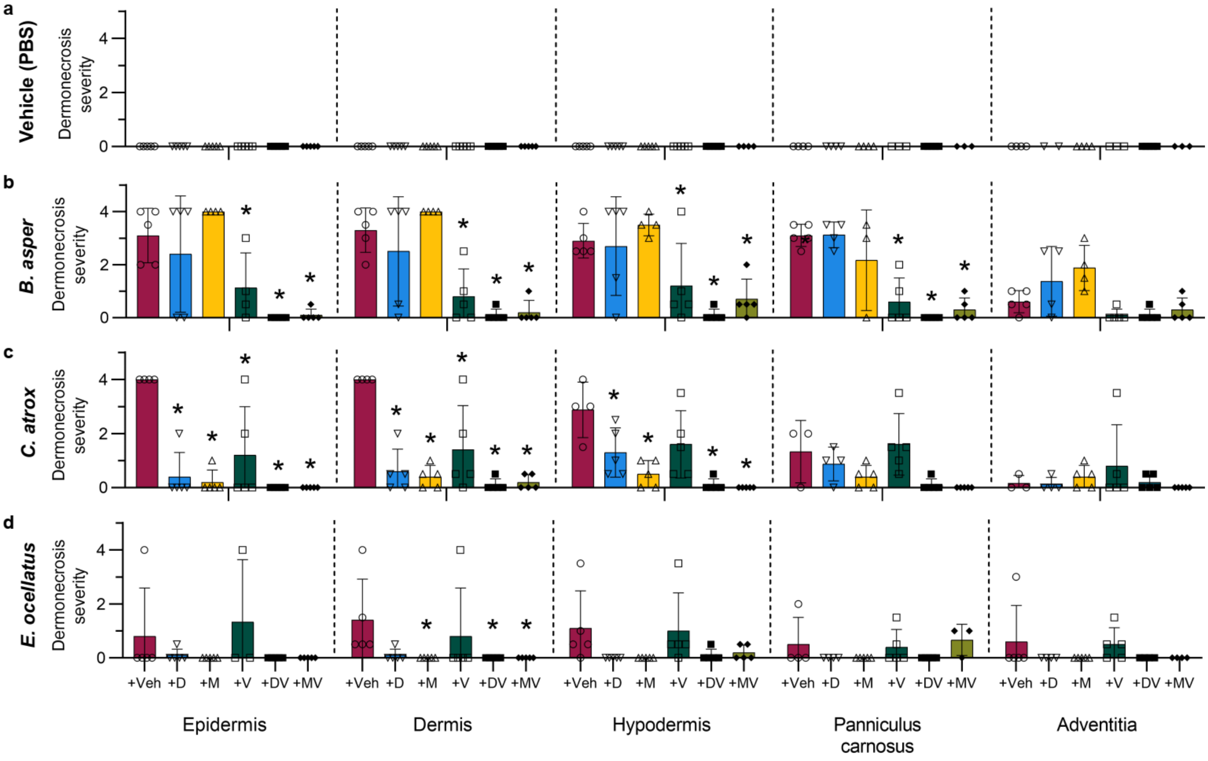


**Supplementary Fig. 4 Histopathological analysis of individual skin layers of the dermal injection site tissue cross-sections.** Four µm H&E sections were prepared from formalin-fixed, paraffin-embedded tissue from dermal lesions and photographed at 100X magnification. Two blinded and independent experimenters scored, between 0-4, the percentage of each skin layer that was necrotic (0 = 0%, 1 = 0-25%, 2 = 25-50%, 3 = 50-75%, and 4 = 75-100% of visible tissue in the image). The highest recorded score per cross-section was used as a measure of the maximum necrotic severity reached within each skin layer for mice ID-injected with (**a**) venom vehicle control (PBS), (**b**) *B. asper* venom (150 µg), (**c**) *C. atrox* venom (100 µg), or (**d**) *E. ocellatus* venom (39 µg) that had been pre-incubated with drug vehicle control (98.48% PBS, 1.52% DMSO; Veh), DMPS (110 µg; D), marimastat (60 µg; M), varespladib (19 µg; V), DMPS-plus-varespladib (110 and 19 µg, respectively; DV), or marimastat-plus-varespladib (60 and 19 µg, respectively; MV), from which mean overall dermonecrosis was determined (**Fig. 6**). * Signifies that value is significantly different than that of the drug vehicle control for that data set as determined by a two-way ANOVA followed by Dunnett's multiple comparisons test ($P < 0.05$, $n \geq 4$). ANOVA statistics can be found in **Supplementary Table 1**. Error bars represent SD of at least 4 scores, and the individual scores are shown as points within each of the figures' bars.

**Supplementary Table 1. Two-Way ANOVA F-values, degrees of freedom, and *P*-values for graphs in Supplementary Fig. 4.**

| Figure letter | Independent variable | F (DFn, DFd) | P value |
|---|---|---|---|
| e | Interaction | N/A* | N/A* |
| | Row factor (skin layer) | N/A* | N/A* |
| | Column factor (drug) | N/A* | N/A* |
| f | Interaction | F (20, 111) = 1.548 | P=0.0795 |
| | Row factor (skin layer) | F (4, 111) = 6.651 | P<0.0001 |
| | Column factor (drug) | F (5, 111) = 41.92 | P<0.0001 |
| g | Interaction | F (20, 111) = 3.112 | P<0.0001 |
| | Row factor (skin layer) | F (4, 111) = 5.299 | P=0.0006 |
| | Column factor (drug) | F (5, 111) = 29.45 | P<0.0001 |
| h | Interaction | F (20, 111) = 0.3648 | P=0.9941 |
| | Row factor (skin layer) | F (4, 111) = 0.3531 | P=0.8414 |
| | Column factor (drug) | F (5, 111) = 5.723 | P<0.0001 |
